# GeneRAG: Enhancing Large Language Models with Gene-Related Task by Retrieval-Augmented Generation

**DOI:** 10.1101/2024.06.24.600176

**Authors:** Xinyi Lin, Gelei Deng, Yuekang Li, Jingquan Ge, Joshua Wing Kei Ho, Yi Liu

## Abstract

Large Language Models (LLMs) like GPT-4 have revolutionized natural language processing and are used in gene analysis, but their gene knowledge is incomplete. Fine-tuning LLMs with external data is costly and resource-intensive. Retrieval-Augmented Generation (RAG) integrates relevant external information dynamically. We introduce GeneRAG, a frame-work that enhances LLMs’ gene-related capabilities using RAG and the Maximal Marginal Relevance (MMR) algorithm. Evaluations with datasets from the National Center for Biotechnology Information (NCBI) show that GeneRAG outperforms GPT-3.5 and GPT-4, with a 39% improvement in answering gene questions, a 43% performance increase in cell type annotation, and a 0.25 decrease in error rates for gene interaction prediction. These results highlight GeneRAG’s potential to bridge a critical gap in LLM capabilities for more effective applications in genetics.

## 1 Introduction

Large Language Models (LLMs) like GPT-4 (Jin et al., 2024) have revolutionized natural language processing (NLP) by enabling highly sophisticated text generation and understanding. Beyond their primary usage in text processing and generation tasks, they have also been actively adopted to solve scientific problems in other areas, including gene-oriented analysis (Hou and Ji, 2024; Xiao et al., 2024; Peng et al., 2024). As shown in the Figure 1, when using LLMs for gene-related analysis, having accurate and up-to-date knowledge about genes significantly influences the precision of the analysis and the validity of its biological conclusions.

**Figure 1:**
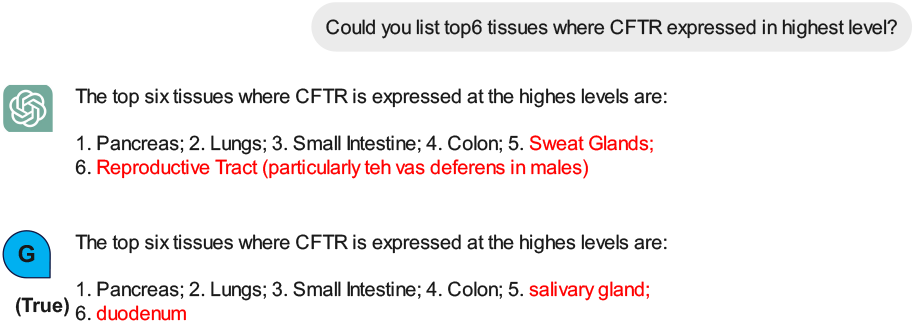
Inaccurate Gene-Related Answers from LLMs.

While LLMs excel in general language tasks, their knowledge about genes is incomplete, and their ability to accurately answer gene-related questions remains limited. This limitation is primarily due to the vast and intricate nature of genetic knowledge, which is difficult to encapsulate within the pre-trained parameters of an LLM. Despite this flaw, the high computational resources and financial costs required make it challenging to fine-tune existing LLMs to enhance them with external knowledge bases. Existing approaches typically investigate the performance of LLMs in various tasks, demonstrating their strengths and weaknesses to guide users in using LLMs properly (Hu et al., 2023; Park et al., 2023). While this is useful, it does not solve the underlying problem. Alternatively, some efforts focus on training new models directly using gene information (Hwang et al., 2024), which is time- and cost-intensive and still struggles to seamlessly integrate extensive and dynamic external knowledge.

Retrieval-Augmented Generation (RAG) represents a promising solution by combining the generative capabilities of LLMs with the precision of retrieval-based methods. RAG dynamically retrieves relevant information from external databases, integrating this knowledge into the generation process to produce more accurate and contextually appropriate responses. This approach is particularly well-suited for addressing the complexities of gene-related inquiries.

In this work, we aim to enhance LLMs’ ability to answer gene-related problems by effectively bridging external knowledge bases through the use of RAG. Our study focuses on addressing the following research questions: (1) How effectively does GeneRAG answer gene-related questions? (2) How effectively does GeneRAG perform gene-oriented downstream analysis?

Our contributions are threefold: First, we propose a novel framework, GeneRAG (available here (Gen)), for integrating external genetic knowledge into LLMs via RAG. Second, we enhance retrieval quality using the Maximal Marginal Relevance (MMR) algorithm. Third, we conduct a comprehensive evaluation of GeneRAG to assess its effectiveness. Using datasets from the National Center for Biotechnology Information (NCBI) and other reliable sources, we created Q&A datasets and selected cell types and genes for downstream tasks. The results show that GeneRAG significantly outperforms GPT-3.5 and GPT-4, achieving higher accuracy and lower error rates. On average, GeneRAG demonstrates a 39% improvement over GPT-4 in answering gene-related questions, a 43% performance increase in cell type annotation, and a 0.25 decrease in error rates for gene interaction prediction. These findings highlight GeneRAG’s potential as a superior tool for both answering gene-related queries and performing downstream gene analysis. The significance of our work lies in its potential to bridge a critical gap in LLM capabilities, enabling more effective and informed applications in genetics and beyond.

## 2 Related Work

### 2.1 LLMs in Gene-Oriented Analysis

Pioneering work in applying language models to gene-oriented analysis falls into two main areas. The first involves directly using LLMs and evaluating their performance. In gene set analysis, LLMs are guided by prompts to identify biological processes and functions of gene groups (Hu et al., 2023; Joachimiak et al., 2023) and their interactions (Azam et al., 2024), with accuracies around 0.5. In cell annotation, GPT-4 predicts cell types based on the top 20 highly expressed genes, achieving up to 0.85 accuracy for well-understood tissues and around 0.2 for less common tissues like the small intestine (Hou and Ji, 2024).

Recognizing limitations in direct usage, researchers have enhanced LLMs’ gene understanding. GeneGPT increased performance to 0.83 by using NCBI Web APIs (Jin et al., 2024). Levine et al. transformed gene expression data into “cell sentences” to fine-tune GPT-2 (Levine et al., 2023). These methods rely on LLMs having up-to-date and accurate knowledge.

### 2.2 RAG

RAG is a hybrid approach that combines the generative capabilities of LLMs with the precision of retrieval-based methods. RAG retrieves relevant information from external knowledge bases and integrates it into the generation process, resulting in more accurate and contextually relevant responses. This method leverages both the vastness of LLMs’ training data and the specificity of up-to-date external knowledge, making it particularly effective for domain-specific tasks.

## 3 Methodology

### 3.1 Overview

The workflow of GeneRAG, depicted in Figure 2, involves a multi-step process designed to enhance the accuracy of answers to gene-related queries. It begins with extracting detailed gene information from an external gene database. This extracted data is then processed by a LLM to create database embeddings, forming a vector database. When a user inputs a prompt, GeneRAG utilizes these embeddings to detect similarities between the prompt and the database entries. By incorporating retrievalaugmented generation, GeneRAG leverages these similarities to provide improved and accurate answers, thus enhancing the overall response quality.

**Figure 2:**
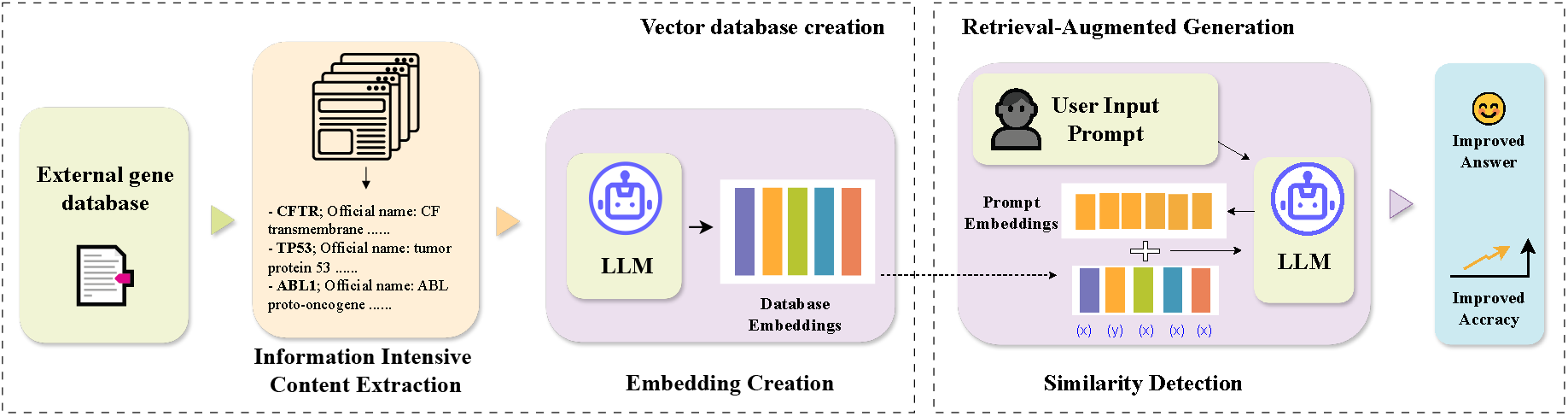
The workflow of GeneRAG.

### 3.2 Data Extraction

We start by collecting gene information from reliable external databases. Specifically, we use data from NCBI (Sherry et al., 2001), which provides comprehensive and up-to-date gene information. This data includes gene names, functions, expressions, and related biological processes. We chose NCBI due to its credibility and the depth of its gene-related data.

To prepare the data for our model, we preprocess it by normalizing the text, removing duplicates, and ensuring consistency in gene terminology. This preprocessing ensures that the information fed into the LLM is clean and standardized, which is crucial for accurate embedding creation.

### 3.3 Embedding Creation

Once we have the processed gene data, we use a LLM to create embeddings. We chose an LLM like *text-embedding-3-large* (Harris et al., 2024) because of its ability to understand and generate human-like text, making it suitable for creating meaningful embeddings from the gene information.

These embeddings are vector representations of the gene data, capturing the semantic meaning of the information. We create a vector database from these embeddings, which allows us to efficiently search and compare the data later. This step is crucial as it forms the foundation for matching user queries with the relevant gene information.

### 3.4 Prompt Handling

When a user inputs a prompt, GeneRAG processes it to determine its intent and content. The system converts the prompt into an embedding using the embedding LLM to facilitate accurate similarity detection.

For detecting similarity, we use cosine similarity. Cosine similarity measures the cosine of the angle between two vectors, which in our case are the embeddings of the prompt and the gene data. It is a widely used method in natural language processing due to its effectiveness in capturing semantic similarity between texts. By using cosine similarity, we can accurately match user queries with the most relevant gene information in our database.

The rationale for using embeddings here is that they provide a powerful way to represent the semantic meaning of the text, enabling the system to understand and match complex queries effectively. By detecting similarities between the prompt embedding and the database embeddings, GeneRAG can identify the most relevant gene information.

### 3.5 Retrieval-Augmented Generation

In the final step, GeneRAG uses retrieval-augmented generation to provide accurate and contextually relevant answers. We employ the Maximal Marginal Relevance (MMR) algorithm to enhance the retrieval process. MMR balances relevance and diversity in the retrieved results, ensuring that the information is both pertinent and non-redundant.

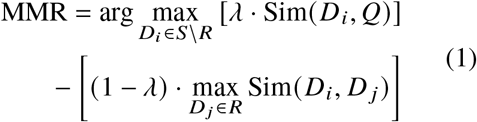

where *D*_*i*_ is a candidate document, *R* is the set of selected documents, *Q* is the query embedding, Sim(*D*_*i*_, *Q*) is the similarity between document *D*_*i*_ and the query, and Sim(*D*_*i*_, *D* _*j*_) is the similarity between documents.

This approach ensures that the selected documents are relevant to the query while also providing diverse information, leading to more comprehensive and accurate answers. The RAG prompt is shown in Appendix.

## 4 Evaluation

To evaluate the effectiveness of GeneRAG, we aim to answer the following two research questions:

**RQ1: (Q&A Effectiveness)**: How effectively does GeneRAG answer gene-related questions?

**RQ2: (Downstream Task Effectiveness)**: How effectively does GeneRAG perform gene-oriented downstream analysis?

### 4.1 Dataset

#### RAG Dataset

We collect 20350 Homo sapiens gene information from NCBI as the external knowledge source due to its reliability and up-to-date gene information.

#### Q&A Dataset

We craft a Q&A dataset consisting of random samples from the *RAG Dataset* with 9,000 questions across three question types, 3000 for each, as shown in Table 2.

#### Downstream Task Dataset

We assess the performance of cell type annotation and gene interaction prediction by selecting 3000 cells and genes from two datasets containing samples from whole human bodies (Eraslan et al., 2022; Han et al., 2020).

### 4.2 Downstream Tasks

Beyond the Q&A task, we also evaluate GeneRAG on other gene-oriented downstream tasks using LLMs. Specifically, we select two downstream tasks.

#### Cell Type Annotation

Cell type annotation is a crucial step in gene-oriented analysis. Genes that are highly expressed in cells indicate cell type information. GPTCelltype (Hou and Ji, 2024) uses LLMs to generate cell type annotations based on the highly expressed genes in each cell. Following this method, we selected 3,000 cells and instruct GeneRAG to annotate their cell types.

#### Gene Interaction Prediction

Gene interaction prediction helps us understand how genes interact with each other. We created a dataset containing known interaction relationships between different genes, following previous work (Azam et al., 2024). In this task, we provided the cell type and gene name information and instruct GeneRAG to list other genes that interact with the query gene.

### 4.3 Metrics

We use three metrics to gauge the effectiveness of GeneRAG. A detailed discussion about the selection of these metrics can be found in Appendix A.

#### Accuracy

Accuracy measures the proportion of correct predictions made by GeneRAG out of all predictions. It is a common metric for evaluating overall performance.

#### False Positive Rate (FPR)

It indicates the proportion of incorrect positive predictions among all actual negative instances. It helps in understanding the rate at which GeneRAG incorrectly identifies non-relevant information as relevant.

#### False Negative Rate (FNR)

This measures the proportion of incorrect negative predictions among all actual positive instances. It shows how often GeneRAG misses relevant information.

### 4.4 RQ1: Q&A Effectiveness

Based on the results presented in Table 1, GeneRAG consistently outperforms both GPT-3.5 and GPT-4o across all question types in terms of accuracy and error rates. On average, GeneRAG demonstrates a 39% improvement compared to GPT-4o. Notably, there is a significant decrease in the false negative rate for trap questions, reduced by 65%. This improvement is likely due to GeneRAG’s reliable reference information, enabling it to focus on facts and ignore traps in the questions. Additionally, GeneRAG exhibits a 43% increase in accuracy for exact answer questions, attributed to its access to trustworthy data sources. These results suggest that GeneRAG is more effective and reliable for answering gene-related questions, establishing it as a superior tool for downstream analysis.

**Table 1:** Performance of GPT and GeneRAG.

| Task | Metric | GPT-3.5 | GPT-4o | GENERAG |
| --- | --- | --- | --- | --- |
| Q&A: Exact Answer | Accuracy | 0.504 ± 0.03 | 0.574 ± 0.04 | 0.824 ± 0.05 |
| Q&A: Multiple Answers | Accuracy | 0.587 ± 0.04 | 0.640 ± 0.06 | 0.763 ± 0.04 |
|  | FPR | 0.211 ± 0.07 | 0.189 ± 0.03 | 0.129 ± 0.02 |
|  | FNR | 0.213 ± 0.07 | 0.180 ± 0.03 | 0.127 ± 0.02 |
| Q&A: Trap Question | FNR | 0.471 ± 0.04 | 0.433 ± 0.05 | 0.151 ± 0.04 |
| Cell Type Annotation | Scores | 0.501 ± 0.04 | 0.594 ± 0.03 | 0.839 ± 0.02 |
| Gene Interaction Prediction | FNR | 0.543 ± 0.05 | 0.576 ± 0.04 | 0.323 ± 0.05 |
|  | Accuracy | 0.594 ± 0.03 | 0.722 ± 0.02 | 0.815 ± 0.03 |

**Table 2:** Questions and metrics of RQ1.

| Question Type | Example | Metrics |
| --- | --- | --- |
| Exact Answer | What is the official name of [gene]? | Accuracy |
| Multiple Answers | List top 6 tissues where [gene] is expressed at highest levels? | Accuracy, FPR, FNR |
| Trap Question | Which of the following functions is [gene] not involved in? | FNR |

FPR: False positive rate. FNR: False negative rate.

### 4.5 RQ2: Downstream Task Effectiveness

In downstream analysis tasks, GeneRAG consistently surpasses GPT-3.5 and GPT-4, as shown in Table 1. In the cell type annotation task, GeneRAG addresses the challenge of insufficient information for rare cell types by utilizing external knowledge, achieving significant improvements over GPT-3.5 and GPT-4 (66% and 41% increases, respectively). Verifying gene-gene interactions with reliable sources also enhances accuracy and reduces error rates. Notably, the false negative rate does not significantly decrease with GPT-4o compared to GPT-3.5, indicating that merely increasing inference capability and training on more data with low-quality information does not substantially improve performance. However, applying high-quality external data proves to be useful.

## 5 Conclusion

In this paper, we introduced GeneRAG, a frame-work for enhancing LLMs in gene-related tasks using RAG. By leveraging the MMR algorithm, GeneRAG improves retrieval quality and provides more accurate responses. Evaluations show that GeneRAG outperforms GPT-3.5 and GPT-4 in gene-related questions, cell type annotation, and gene interaction prediction. These findings highlight GeneRAG’s potential to bridge the gap between LLMs and external knowledge bases, advancing their application in genetics and other scientific fields.

## 6 Limitations

Despite the promising results, our study has several limitations. First, the reliance on external knowledge bases means that the performance of GeneRAG is heavily dependent on the quality and completeness of these sources. Any inaccuracies or gaps in the external databases can directly affect the output of GeneRAG. Second, while GeneRAG shows significant improvements in specific tasks, its effectiveness may vary across different types of gene-related questions and downstream tasks not covered in this study. Third, the computational overhead associated with the retrieval-augmented generation approach can be substantial, potentially limiting its scalability and practical application in resource-constrained environments. Finally, our evaluation datasets, although comprehensive, may not fully capture the diversity and complexity of real-world genetic data, which could affect the generalizability of our findings.

## 7 Ethical Statement

In conducting this research, we have adhered to the highest ethical standards. All data used in this study were obtained from publicly available and reputable sources, such as NCBI, ensuring that no private or sensitive information was used. Our study aims to advance scientific knowledge and improve gene-related analysis through the development of GeneRAG. We recognize the potential implications of using large language models and strive to ensure that our work promotes positive and responsible use of artificial intelligence in scientific research. Additionally, we have taken care to validate our findings thoroughly to prevent the dissemination of inaccurate or misleading information. Any potential biases and limitations of our approach have been transparently reported to facilitate further research and improvement in this field.

## A Details explanation of evaluation metrics

For RQ1, we designed three metrics to evaluate the performance for three question types, as shown in Table 2. For **Exact Answer** questions, there is only one correct answer, and the performance is described by accuracy. For **Multiple Answers** questions, like *“List top 6 tissues where [gene] is expressed at highest levels?”*, the correct answers are 6 tissues. Missing tissues are marked as false negatives, while wrong tissues are marked as false positives. The metrics for evaluation include accuracy, false positive rate (FPR), and false negative rate (FNR). **Trap Questions**, exemplified by *“Which of the following functions is [gene] not involved in?”*, are evaluated using the false positive rate because all listed functions should involve the gene, so the answer to this question should be ‘none of them’.

For RQ2, the performance in the cell type annotation task is evaluated by a score where 1, 0.5, and 0 indicate ‘fully match’, ‘partially match’, and ‘mismatch’, respectively. For the gene interaction prediction task, we first use accuracy to describe the number of correctly predicted genes and use the false negative rate to describe the proportion of genes the tool predicts as having no interaction relationship when, in fact, there is an interaction relationship.

## B RAG prompt

#### RAG Prompt

You are an assistant for question-answering tasks. Use the following pieces of retrieved context to answer the question. If you don’t know the answer, just say that you don’t know. Use three sentences maximum and keep the answer concise.

Question: {question}

Context: {context}

Answer:

## C Description of data

### NCBI

The National Center for Biotechnology Information (NCBI) is a key resource in the field of biotechnology and bioinformatics, providing access to a vast array of databases and tools for the scientific community. Established in 1988 as part of the National Library of Medicine (NLM) at the National Institutes of Health (NIH), NCBI is dedicated to the organization and dissemination of biomedical information. The Gene database in NCBI integrates information about genetic sequences, including nomenclature, chromosomal localization, gene products, and phenotypic effects.

### Downstream dataset1

This study utilized microwell-seq, a cost-effective single-cell mRNA sequencing technology, to construct a comprehensive Human Cell Landscape (HCL) by profiling 702,968 single cells across 60 human tissue types, including both fetal and adult samples from a Chinese Han population. The analysis, which covered approximately 3,000 reads per cell, revealed 102 major cell-type clusters and 843 sub-clusters, demonstrating high gene and cell-type coverage. Utilizing high-throughput barcoding strategies, the study achieved extensive mapping of cell types, including stromal/mesenchymal cells, endothelial cells, macrophages, and fetal epithelial cells. The t-SNE analysis visualized 599,926 single cells, identifying 102 distinct clusters with multi-tissue contributions. Further, the study indicated limited donor or batch effects on cell-type discovery and employed a pseudo-cell concept to enhance gene representation and cluster separation. The resource, representing the most comprehensive cell-type repertoire yet described for humans, is publicly available at bis.zju.edu.cn/HCL/ and db.cngb.org/HCL/. (Han et al., 2020)

### Downstream dataset2

This dataset comprises 209,126 high-quality single-nucleus RNA sequencing (snRNA-seq) profiles from 25 archived, frozen tissue samples obtained from the Genotype-Tissue Expression (GTEx) project. The samples span eight tissue sites—breast, esophagus mucosa, esophagus muscularis, heart, lung, prostate, skeletal muscle, and skin—from 16 individuals (seven males and nine females). Using four distinct nucleus isolation protocols, the nuclei were processed through droplet-based single-cell RNA-seq, detecting an average of 918 genes and 1519 unique molecular identifiers per profile. Corrected for ambient RNA contamination, this dataset provides a comprehensive reference atlas, highlighting the cellular diversity and heterogeneity across multiple tissues and individuals. (Eraslan et al., 2022)

